# MetaCompare 2.0: Differential ranking of ecological and human health resistome risks

**DOI:** 10.1101/2024.01.17.576132

**Authors:** Monjura Afrin Rumi, Min Oh, Benjamin C. Davis, Adheesh Juvekar, Connor L. Brown, Peter J. Vikesland, Amy Pruden, Liqing Zhang

## Abstract

**Background:** While there is increasing recognition of numerous environmental contributions to the spread of antibiotic resistance, quantifying the relative contributions of various sources remains a fundamental challenge. Similarly, there is a need to differentiate acute human health risks corresponding to exposure to a given environment, versus broader ecological risk of evolution and spread of antibiotic resistance genes (ARGs) across microbial taxa. Recent studies have proposed various methods of harnessing the rich information housed by metagenomic data for achieving such aims. Here, we introduce MetaCompare 2.0, which improves upon the original MetaCompare pipeline by differentiating indicators of human health resistome risk (i.e., potential for human pathogens to acquire ARGs) from ecological resistome risk (i.e., overall mobility of ARGs across a given microbiome).

**Results:** To demonstrate the sensitivity of the MetaCompare 2.0 pipeline, we analyzed publicly available metagenomes representing a broad array of environments, including wastewater, surface water, soil, sediment, and human gut. We also assessed the effect of sequence assembly methods on the risk scores. We further evaluated the robustness of the pipeline to sequencing depth, contig count, and metagenomic library coverage bias through comparative analysis of a range of subsamples extracted from a set of deeply sequenced wastewater metagenomes. The analysis utilizing samples from different environments demonstrated that MetaCompare 2.0 consistently produces lower risk scores for environments with little human influence and higher risk scores for human contaminated environments affected by pollution or other stressors. We found that the ranks of risk scores were not measurably affected by different assemblers employed. The MetaCompare 2.0 risk scores were remarkably consistent despite varying sequencing depth, contig count, and coverage.

**Conclusion:** MetaCompare 2.0 successfully ranked a wide array of environments according to both human health and ecological resistome risks, with both scores being strongly impacted by anthropogenic stress. We packaged the improved pipeline into a publicly-available web service that provides an easy-to-use interface for computing resistome risk scores and visualizing results. The web service is available at http://metacompare.cs.vt.edu/

## 1 Introduction

Antibiotic resistance is a global public health threat, resulting in an increasing rate of human morbidity and mortality worldwide [1]. There are numerous sources, pathways, and factors that contribute to the evolution and spread of antibiotic resistance, which makes it difficult to pinpoint precise interventions that can help attenuate the carriage of antibiotic resistance genes (ARGs) by human pathogens. It is increasingly being recognized that mitigation efforts must move beyond a myopic focus on clinical settings and must address environmental sources and ecological processes that contribute to the spread of resistance [2, 3]. Environmental sources of concern include untreated sewage, wastewater treatment plant (WWTP) effluent, livestock waste, surface water runoff, landfill leachate, and pharmaceutical manufacturing waste [4].

To effectively inform strategies to mitigate the spread of antibiotic resistance, a systematic and quantitative means of comparing putative sources of ARGs and their potential to be acquired by pathogens is required [5, 6]. For this purpose, MetaCompare [7], herein referred to as MetaCompare 1.0, was introduced as the first computational pipeline to quantify and rank the “resistome risk” of various environments. “Resistome risk” refers to the conceptual framework introduced by Martínez et al. [6], in which it is assumed that ARGs that 1) confer resistance to antibiotics currently used for therapeutic purposes, 2) are associated with mobile genetic elements (MGEs), and 3) are carried by human pathogens represent the greatest public health risk. MetaCompare 1.0 was developed as a pipeline to put this concept into practice and introduced a reified resistome risk metric [7]. Comparing the resistome risk metric across environments can serve as a means to identify potential “hot spots” for mobilization of antibiotic resistance to pathogens, which can then be prioritized for targeted mitigation. A proof-of-concept experiment using publicly-available metagenomic datasets demonstrated that MetaCompare 1.0 provided a ranking of resistome risk consistent with expectations. Specifically, the pipeline ranked resistomes in order of hospital sewage as having the highest risk scores, dairy lagoons as having moderate risk scores, and WWTP effluent as having the lowest risk scores [7]. MetaCompare 1.0 has now been widely applied, providing insight into potential critical control points for ARG transmission to pathogens across a wide variety of agricultural, wastewater, and other environmental systems e.g., [8–14].

Similar methods have been proposed to assess and rank the risks of individual ARGs in various environments. For example, Slizovskiy et al. [15] proposed a metric called the ‘mobility index’ which considers ARGs being carried by MGEs in a sample, but does not take into account the presence of pathogens. Zhang A.N. et al. [16] developed a method to rank ARGs in terms of their anthropogenic enrichment, association with MGEs (mobility), and human pathogens (host pathogenicity) into four categories, with Rank I being the highest risk category and Rank IV being the lowest. Subsequently, Zhang Z. et al. [17] defined a “risk index” to categorize ARGs prevalent only in human-associated environments considering the clinical availability of antibiotics, mobility, host pathogenicity, and potential of transmission of ARGs from environment to humans. A key distinction relative to other such ARG risk ranking systems is that MetaCompare resistome risk scores are defined for the entire collection of ARGs detected in a sample, whereas the latter provides a ranking system only for individual ARGs. Incorporation of individual ARG risks into the broader resistome risk of the environment represented by a sample remains a distinct computational endeavor.

While it is widely recognized that a risk assessment framework is needed to address environmental dimensions of antibiotic resistance [3], a challenge is that it does not fit the mold of conventional microbial risk assessment [18]. Specifically, there are multiple bacterial pathogens of concern and thousands of ARGs. Considering exposures to individual resistant pathogens can inform quantitative microbial risk assessment [19, 20], but evolution and horizontal gene transfer of ARGs moving across microbial communities, including both pathogens and non-pathogens, is arguably of equal concern if the aim is to mitigate the acquisition of ARGs by pathogens in the first place. Here we use the term “risk” broadly as a general relative comparison, as frameworks remain to be adapted to move towards estimating probabilities of resistant infections from dose-response of various resistome exposures. From a human health risk standpoint, the ESKAPE pathogens (*Enterococcus faecium, Staphylococcus aureus, Klebsiella pneumoniae, Acinetobacter baumannii, Pseudomonas aeruginosa, and Enterobacter spp*.), have been recognized as World Health Organization (WHO) priority pathogens that tend to be highly virulent and antibiotic resistant [21]. ESKAPE pathogens have also been shown to be enriched with a specific subset of acquired (i.e., not belonging to their core genome), mobile ARGs in anthropogenically-impacted environments [16]. From an ecological standpoint, a major limitation has been a lack of a suitable database for accurate annotation of MGEs. Inclusion of accessory or cargo sequences, including ARGs, in public MGE databases, such as ACLAME [22], has undoubtedly led to false positives (e.g., an ARG being annotated as an MGE) [15]. Finally, it is not clear how differing sequencing and analysis approaches affect the risk scores. In the case of MetaCompare 1.0, *de novo* assembly of contigs is required, the representativeness of which is directly affected by sequencing depth, chosen assembler, and microbial diversity associated with the sample complexity.

Here we introduce MetaCompare 2.0, which incorporates several improvements to address the above-noted limitations of MetaCompare 1.0 and other ARG risk ranking approaches. Specifically, we introduce two distinct resistome risk scores, one corresponding to the ecological resistome risk (ERR) and the other to the human health resistome risk (HHRR). The ERR score factors in a wide-ranging array of pathogens and ARGs in order to broadly represent the potential for ARGs to mobilize in a given environment. The HHRR, on the other hand, focuses more specifically on pathogens that are of most acute concern with regard to antibiotic resistant infections in humans (i.e., ESKAPE pathogens) and Rank I ARGs [16]. For both indices, the annotation methodology for the taxonomic assignment of potential pathogens was improved by using ‘many-against-many’ sequence searching (MMseqs2), a tool designed specifically for taxonomy assignment [23]. Inaccuracies in MGE annotation were addressed by incorporating an updated MGE database (mobileOG-DB) [24]. To provide a more intuitive output for comparison, the range of possible values for risk scores was set using a 0-100 scale. In the case of the ERR specifically, DeepARG database (DeepARG-DB) [25], a database built specifically to capture environmental ARGs, was applied to broadly consider the potential for antibiotic resistance to evolve and spread. To assess an expanded range of feasible output resistome risk values, the updated pipeline was applied to publicly-available metagenomes representing a wide range of environments, including wastewater, surface water, soil, sediment, and gut microbiomes. MetaCompare 2.0 was further validated by applying on samples having varying sequencing depths, assembler methods, and assembly sizes. Lastly, the pipeline was made more accessible through a web service that allows users to compute the risk scores of their metagenomic data and visualize annotations of assembled sequences in various dimensions.

## 2 Methods

### 2.1 Overview of pipeline

MetaCompare 2.0 employs two computational branches that can be selected by the user (Figure 1). One branch assesses the ERR and the other assesses the HHRR. The ERR evaluates a broad array of both known and putative ARGs, their co-occurrence with MGEs, and a full range of human bacterial pathogens to capture their probable contribution to the proliferation of antibiotic resistance in corresponding environments. The HHRR focuses on a narrower set of ARGs, defined by [16] as Rank I ARGs, that are: 1) demonstrated to be enriched in “human-associated” environments, 2) mobile (carried by MGE), and 3) can be carried by ESKAPE pathogens. The range of pathogens used in HHRR was confined to the ESKAPE pathogens as well as any contig annotated as Enterobacteraciae. This was done to include Escherichia coli (as in the “ESKAPEE”) [26, 27] as well as to ensure that contigs without complete taxonomic annotations but which are likely derived from the ESKAPE organisms would be captured. This is common for contigs originating from ESKAPE-associated plasmids and MGEs [28, 29].

**Fig. 1.**
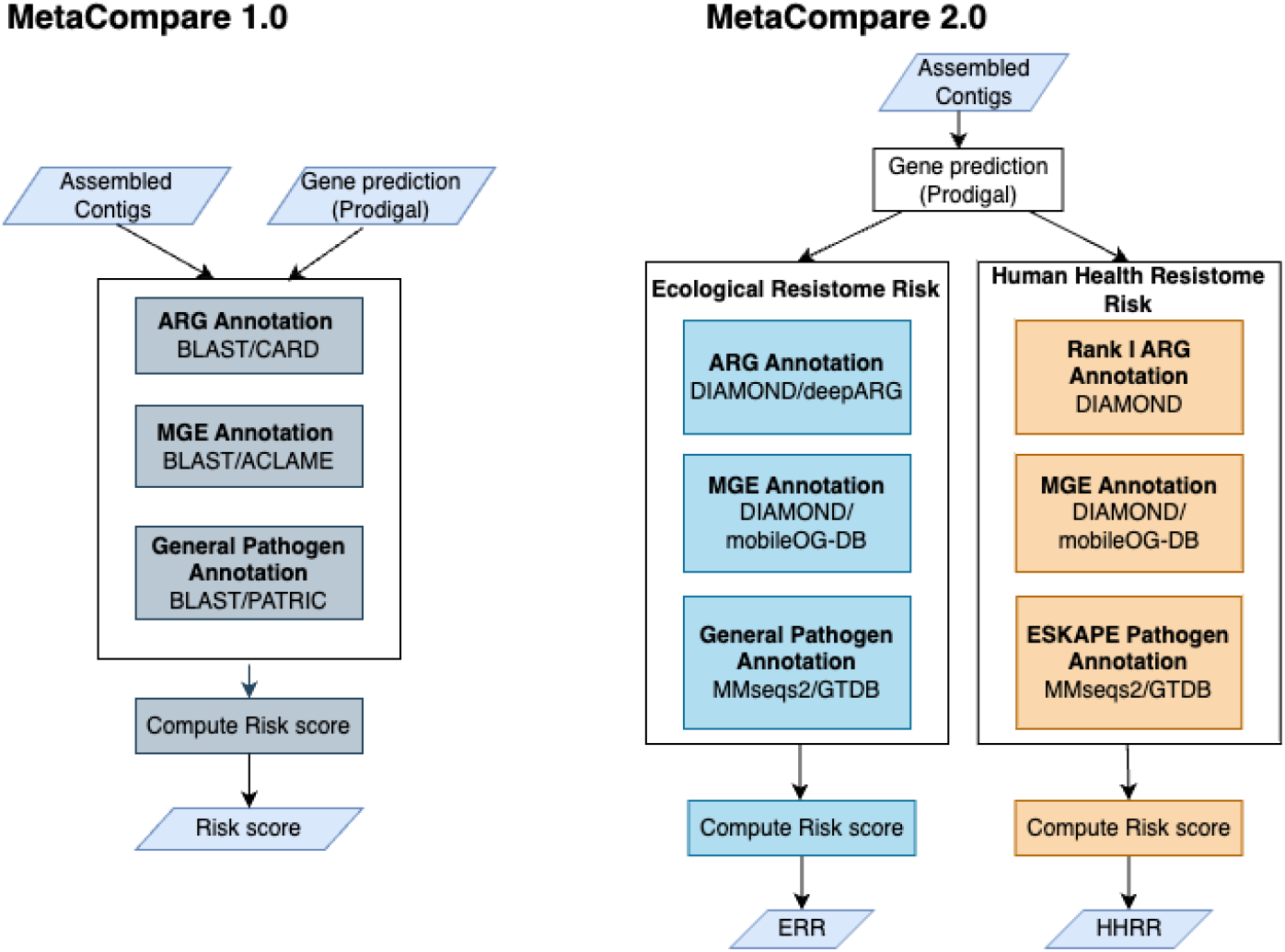
Modifications of the MetaCompare 2.0 pipeline relative to the MetaCompare 1.0 pipeline

MetaCompare 2.0 follows the computational approach adopted in MetaCompare 1.0, with some modifications. MetaCompare 1.0 pipeline takes assembled contigs as input and annotates them in terms of ARGs, MGEs, and taxonomic similarity to known human pathogens. Subsequently, these contigs are classified into three categories: 1) those containing one or more ARGs, 2) those containing one or more ARGs and one or more MGEs, or 3) those containing one or more ARGs, MGEs, and alignment to known human pathogens. The numbers of contigs belonging to these three categories are subsequently normalized by the total number of contigs using Equations 1-3. The normalization is unweighted in terms of the number of ARGs or MGEs co-occurring on a contig. Since a pathogen with an ARG is still a human health concern, even if it cannot be demonstrated mobile, a new category has been added in MetaCompare 2.0 for contigs containing one or more ARGs and alignment to known human pathogens (Equation 4).

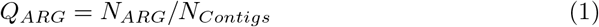

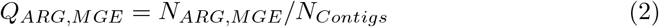

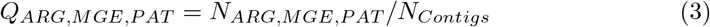

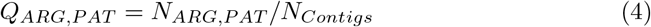

where *N*_*Contigs*_ is the total number of contigs in the sample and *N*_*ARG*_, *N*_*ARG,MGE*_, *N*_*ARG,MGE,PAT*_, *N*_*ARG,PAT*_ are the numbers of contigs that contain regions annotated as ARGs only, contain annotated regions indicating that ARGs are proximal to MGEs, contain annotated regions indicating that MGE-associated ARGs are carried within a pathogen, and contain annotated regions indicating that an ARG is carried by a pathogen, respectively.

Additional modifications were introduced to make the risk score output more intuitive. MetaCompare 1.0 calculates the risk score by projecting the samples in a 3-dimensional space termed as ‘hazard space,’ each dimension corresponding to the proportions of contigs annotated as carrying ARGs, carrying ARGs and MGEs, and carrying ARGs, MGEs, and having alignment to known human pathogens, respectively. An empirical theoretical maximum point, indicating the highest value any Q in Equation 1-3 can reach, was set to (0.01, 0.01, 0.01) based on the result of a simulation utilizing the prevalence data in the Comprehensive Antibiotic Resistance Database (CARD) [30]. However, subsequent studies obtained values exceeding this maximum threshold [31]. Therefore we reset the maximum point to (1.0, 1.0, 1.0) to calculate *ds*, the Euclidean distance of the sample to the maximal point in the 3D hazard space with dimensions *Q*_*ARG*_, *Q*_*ARG,MGE*_, *Q*_*ARG,MGE,PAT*_. We also calculated *dw*, the Euclidean distance of a sample containing no ARGs (i.e., *Q*_*ARG*_, *Q*_*ARG,MGE*_, *Q*_*ARG,MGE,PAT*_ are all 0) and used *dw* to normalize *ds*. The basic framework of the 3D hazard space was maintained in the calculation of ERR in MetaCompare 2.0. However, a 4th dimension, *Q*_*ARG,PAT*_, was added for HHRR calculation. The risk score is simplified as follows:

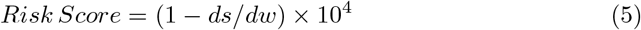

By removing the distance inversion and logarithmic scaling used in MetaCompare 1.0, we have alleviated the problem of producing a theoretically infinite score. Instead, we have set the minimum score to 0, where it was originally 17.57. The maximum score is now set at 100, by incorporating a multiplication factor of 10^4^ (Equation 5).

### 2.2 Updates in tools and databases

Updates incorporated into MetaCompare 2.0 are summarized in Table 1.

**Table 1.**
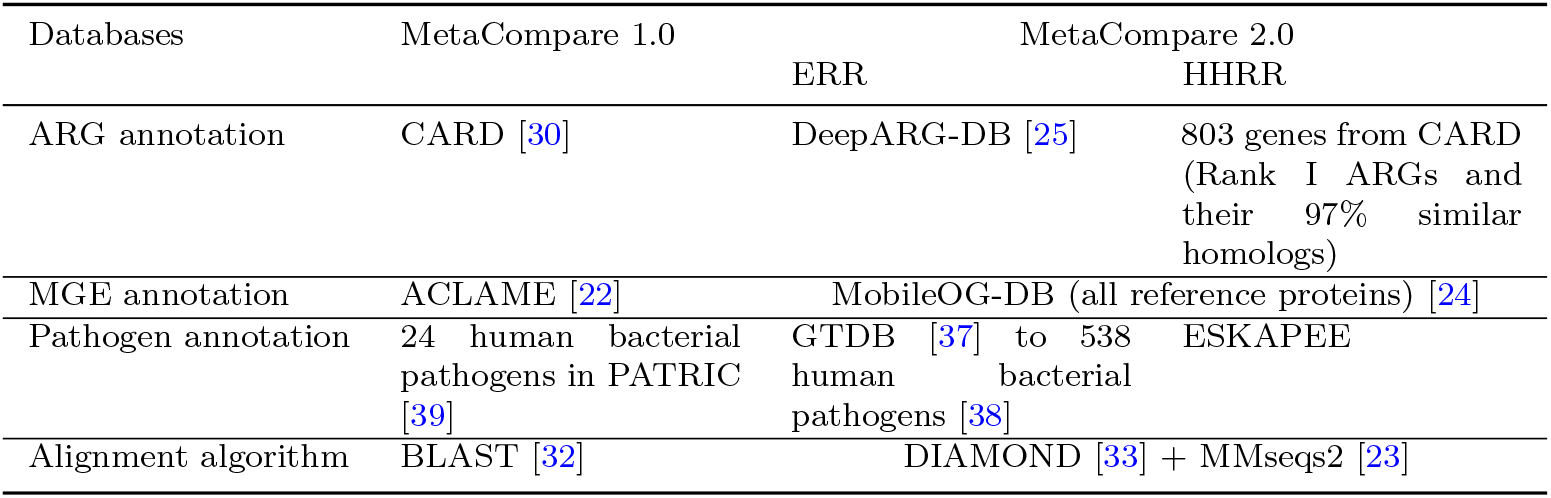
Summary of updates in MetaCompare 2.0 relative to MetaCompare 1.0.

While MetaCompare 1.0 used the standard Basic Local Alignment Search Tool (BLAST)) [32]), MetaCompare 2.0 incorporates DIAMOND BLASTx [33] for ARG and MGE annotation. DIAMOND dramatically improves processing time compared to BLAST through its employment of double indexing for search and alignment, which is more efficient for large environmental microbiome datasets. Prior studies have demonstrated highly comparable consistent results of DIAMOND and BLAST alignment [33, 34]. For pathogen annotation, MMseqs2 [23] replaces BLAST in MetaCompare 2.0. MMseqs2 uses k-mer searches based on similarity instead of exact matches which enables it to use long k-mers without losing sensitivity. Elimination of random memory access and parallelization on multiple levels makes the runtime of MMseq2 highly scalable and thus it has become a popular tool for species/taxonomy annotation [23].

The sensitivity and relevance of annotations have also been improved by updating the associated databases queried by MetaCompare 2.0. DeepARG-DB was employed for ARG annotation because it was constructed using a deep learning algorithm to capture all known and putative ARGs in a metagenome, including ones that may not yet be reported in public databases [25]. DeepARG-DB was built by integrating multiple databases (CARD, Antibiotic Resistance Genes Database [35] and Universal Protein Resource [36]) in a non-redundant fashion. Expanded ARG detection is especially important for the calculation of ERR, where the aim is to assess the potential of antibiotic resistance to evolve and mobilize in a given environment. For MGEs, the mobile orthologous groups database (mobileOG-DB) was incorporated. Extensive manual curation was employed in mobileOG-DB to comprehensively include multiple MGE types (i.e., plasmids, transposons, integrons, etc.) while excluding accessory and cargo genes to avoid false positive annotations [24]. Genome taxonomy database (GTDB) [37] was used to annotate contigs for pathogens. Pathogens were classified against GTDB and filtered from MMseqs2 output using a predetermined list of 538 known, emerging, and re-emerging bacterial pathogens [38]. We selected genus, species, and strain level annotations of MMseqs2 for pathogen filtration. The list of included pathogens is presented in supplementary Table S1. To expand the capabilities of MetaCompare 2.0, we downloaded the list of 122 Rank I ARG references from [16] and aligned them to the protein homolog model database of CARD (v3.2.0) [30] using BLASTp [33]. The alignments were then filtered at *≥* 97% identity to expand the list of Rank I ARGs to include the original set plus their closest homologs, resulting in 803 total ARGs. The expanded list of ARGs included in HHRS calculation is reported in Supplementary Table S2.

### 2.3 Web Service

MetaCompare 2.0 has been made publicly-available as a web service to increase accessibility and ease of use (http://metacompare.cs.vt.edu/). Users can upload assembled metagenomic FASTA files associated with samples of interest and can process the pipeline directly from the web server using a user-friendly graphical interface. A back-end server performs all necessary computational analysis while the front-end service presents the results in tabular format, which can be downloaded as a CSV file. In the command-line interface of MetaCompare 1.0, users were required to provide two FASTA input files: one containing the assembled contigs and the other containing their predicted protein coding regions. In the web platform, users only need to upload the assembled contigs and the burden of computing gene prediction is taken care of in the back-end. Finally, to provide informative output and allow users to inspect further and investigate annotated contigs, a new visualization functionality has been incorporated. Specifically, the visual output allows users to inspect which specific ARGs, MGEs, and pathogens are annotated and their relative positions on the contigs. Researchers can zoom in/out and extract corresponding ARG/MGE/pathogen DNA sequences for further analysis (Figure 2).

**Fig. 2.**
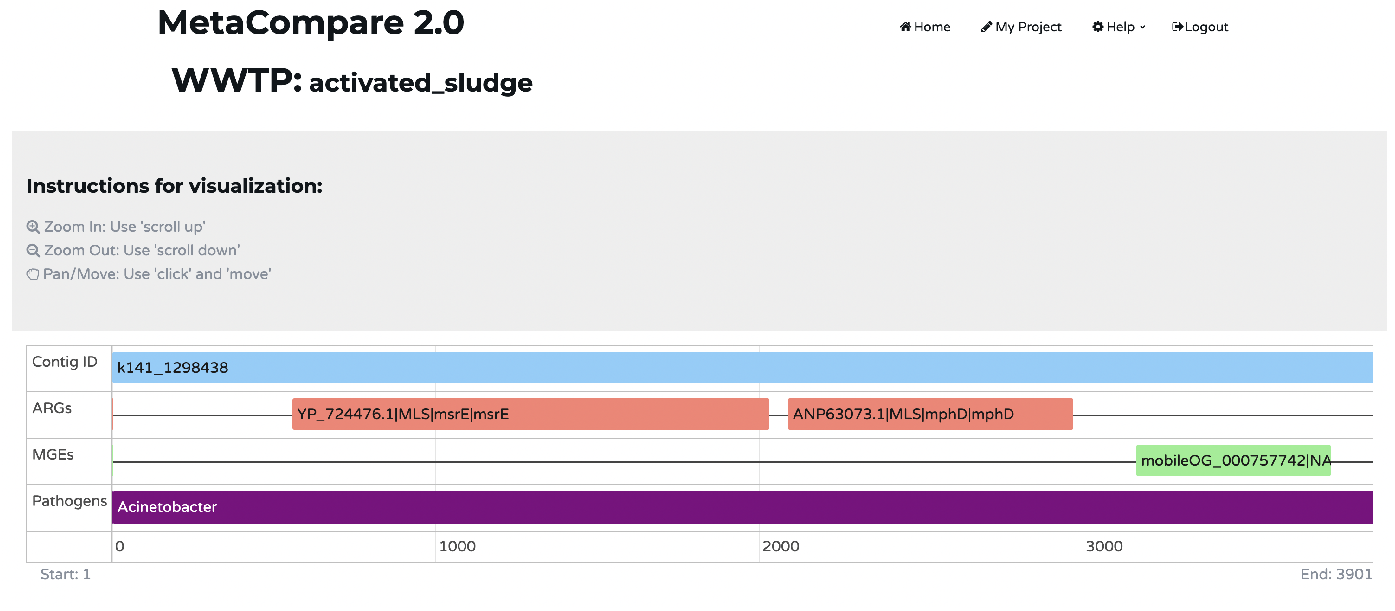
Visualization of ARG, MGE and Pathogen annotation of a representative contig via MetaCompare 2.0 web service.

## 3 MetaCompare 2.0 Validation

Several areas of uncertainty in model robustness were raised during the development of the MetaCompare 1.0 pipeline. Specifically, it was uncertain whether different sample types (i.e., sample complexity), assemblers, library coverage, and the total number of contigs have an effect on the computed risk scores [40, 41]. To challenge both ERR and HHRR models, we collected publicly-available Illumina short reads from NCBI, along with an internal archive of deeply sequenced wastewater metagenomes. The criteria for collection are explained in more detail in section 3.1. All samples were initially cleaned using fastp [42] with default parameters and assembled using MEGAHIT [43], unless otherwise specified. Prodigal [44], a prokaryotic gene recognition tool, was applied for predicting protein-coding genes in contigs.

### 3.1 Survey of diverse environments

The original MetaCompare 1.0 pipeline was tested using three contrasting categories of human influenced aquatic environments: hospital sewage, WWTP effluent, and agricultural lagoon water. Here we used the same dataset for benchmarking MetaCompare 2.0 for initial comparison between the new and old versions. Additionally, we aimed to establish the range of scores encountered when MetaCompare 2.0 was applied to a much wider range of environments, namely: soil, sediment, surface water, gut microbiome, WWTP, as well as mock microbial communities, and deionized water (categorized as “Lab generated” in latter sections) samples. For each environment, we identified contrasting sub-sets of samples based on available SRA metadata and contextual evidence from their accompanying research articles. For each sub-set, we sought to identify 10 samples. We defined the sub-set coming from human influenced environments as a “polluted” set and the sub-set coming from less human influenced environments as “unpolluted”. For example, we refer to samples collected from Arctic soil [45] as “unpolluted” and soil collected from dairy farms (PRJNA379303) as “polluted”. We contrasted sediment samples collected from the river bed of a mountain stream [46] (unpolluted) with river sediments contaminated by pharmaceutical discharge (PRJEB28019) (polluted). For surface water, we compared freshwater samples (PRJNA626373) (unpolluted) with water from ditches (polluted) in densely populated regions (PRJEB13833). For the gut microbiome, we compared samples collected from healthy people and samples from COVID-19 patients [47] since it was found that COVID-19 significantly alters gut microbiome composition [48]. We also tested edge cases (samples with a high probability of generating extremely low/high risk scores) using deionized water (i.e., negative controls) which have a near zero probability of ARG presence and Zymo mock microbial communities (catalog number D6300, zymoresearch.com) which contain mixtures of 10 organisms, 7 of which are bacterial pathogens (*Listeria monocytogenes, Pseudomonas aeruginosa, Bacillus subtilis, Escherichia coli, Salmonella enterica, Enterococcus faecalis, Staphylococcus aureus*). For WWTPs, we contrasted influent (PRJEB13831) and effluent samples (PRJNA438174, PRJNA490743, PRJNA904380, PRJNA505617, PRJEB14051, PRJNA532678, PRJEB15519). For WWTPs, we contrasted raw influent (PRJEB13831) and treated effluent samples (PRJNA438174, PRJNA490743, PRJNA904380, PRJNA505617, PRJEB14051, PRJNA532678, PRJEB15519). All samples, their BioProjects and SRA accessions used in model validation are provided in Supplementary Table S3. MetaCompare 2.0 was applied to determine risk scores of these samples. Wilcoxon rank-sum tests were performed to examine whether the differences between the ERR and HHRR scores of the paired categories were statistically significant.

### 3.2 Evaluation of the Effect of Assembly Method on Resulting Risk Scores

The original MetaCompare 1.0 pipeline used IDBA-UD [49] for short-read assembly. However, it has been demonstrated that assemblers can vary in their accuracy [41], which could potentially affect downstream risk scores. To examine the effect of assembler choices on the risk scores, we applied three commonly used assemblers: IDBA-UD [49], MEGAHIT [43], and metaSpades [50] to five samples from five different environments. The detailed metadata for these samples are provided in Supplementary Table S4. All samples were quality filtered using fastp [42] prior to assembly. For each sample, three sets of contigs were analyzed, one for each assembly approach, prior to MetaCompare 2.0 analysis.

### 3.3 Evaluation of the Effect of Sequencing Depth and Assembly Size

To assess the effect of sequencing depth, coverage, and assembly size on risk scores, we subsampled both short-reads and assembled contigs from deeply sequenced Illumina datasets for this purpose using Seqtk [51]. Short-reads were subsampled from three deeply sequenced wastewater samples that we generated internally from influent, effluent, and activated sludge, each containing *∼* 4 billion reads (*∼* 1TB) sequenced on an Illumina NovaSeq6000. The samples were subsampled at discrete intervals of 1M (million), 10M, 50M, 100M, 125M, and 250M, then assembled with MEGAHIT [43] and scored. Additionally, Nonpareil3 [52] was used on the pre-assembled short-read data to estimate library coverage for each sample using options -T kmer. For contig simulation, we assembled the largest activated sludge sample (250M reads resulting in 6,645,925 contigs) and subsampled sets of contigs containing 2.5k (1k = 1000), 5k, 10k, 50k, 100k, 500k, and 1M contigs, which were then annotated and scored. For each subsampling, we generated 50 sets of contigs.

## 4 Results

The test dataset used to demonstrate the performance of MetaCompare 1.0 was reanalyzed to assess differences in output using MetaCompare 2.0 (Table 2). We assessed normality using the Shapiro-Wilk test and then calculated Pearson’s correlation coefficient between risk scores from MetaCompare 1.0 and MetaCompare 2.0 for this dataset. The Pearson correlation coefficients of ERR and HHRR scores with MetaCompare 1.0 scores were 0.98 (p-value 2.318e−9) and 0.95 (p-value 1.371e−7), respectively. MetaCompare 2.0 more clearly separated dairy lagoon and WWTP effluent samples, relative to MetaCompare 1.0, as highlighted in Table 2.

**Table 2.** MetaCompare 2.0 risk scores obtained from the original MetaCompare 1.0 validation study.

| ENA accession | Sample type | MetaCompare 1.0 | MetaCompare 2.0 |  |
| --- | --- | --- | --- | --- |
|  |  |  | ERR | HHRR |
| ERS1019924 | Hospital sewage | 43.00 | 23.19 | 1.8 |
| ERS1019927 | Hospital sewage | 39.47 | 18.58 | 1.22 |
| ERS1019928 | Hospital sewage | 39.24 | 20.25 | 1.42 |
| ERS1019923 | Hospital sewage | 36.23 | 19.24 | 1.52 |
| ERS1019925 | Hospital sewage | 34.62 | 17.05 | 1.03 |
| ERS1019926 | Hospital sewage | 34.47 | 16.88 | 1.34 |
| ERS1019959 | Dairy lagoon | 29.02 | 10.27 | 1.01 |
| ERS1019958 | Dairy lagoon | 26.84 | 11.15 | 0.5 |
| ERS1019922 | Dairy lagoon | 24.82 | 6.79 | 0.74 |
| ERS1019955 | Dairy lagoon | 24.20 | 6.6 | 0.44 |
| ERS1019956 | **Dairy lagoon** | **22.71** | **7.59** | **0.36** |
| ERS1019920 | **WWTP effluent** | **22.77** | **6.5** | **0.34** |
| ERS1019947 | WWTP effluent | 21.59 | 3.96 | 0.17 |
| ERS1019948 | WWTP effluent | 18.42 | 5.87 | 0.22 |

### 4.1 Range of risk scores encountered across diverse environments

Figure 3 illustrates the range of risk scores encountered across a broad range of environmental metagenomes (e.g., wastewater, surface water, soil, sediment, gut microbiome, and lab generated water). The highest risk score was 63.52, generated by a sample in the Zymo Mock Microbial Community and the lowest score was 0, generated by a deionized water sample. For each environment, the average ERR score and HHRR scores were higher in the “polluted” dataset compared to the “unpolluted” (Wilcoxon rank sum test; p-value *<* 0.0003).

**Fig. 3.**
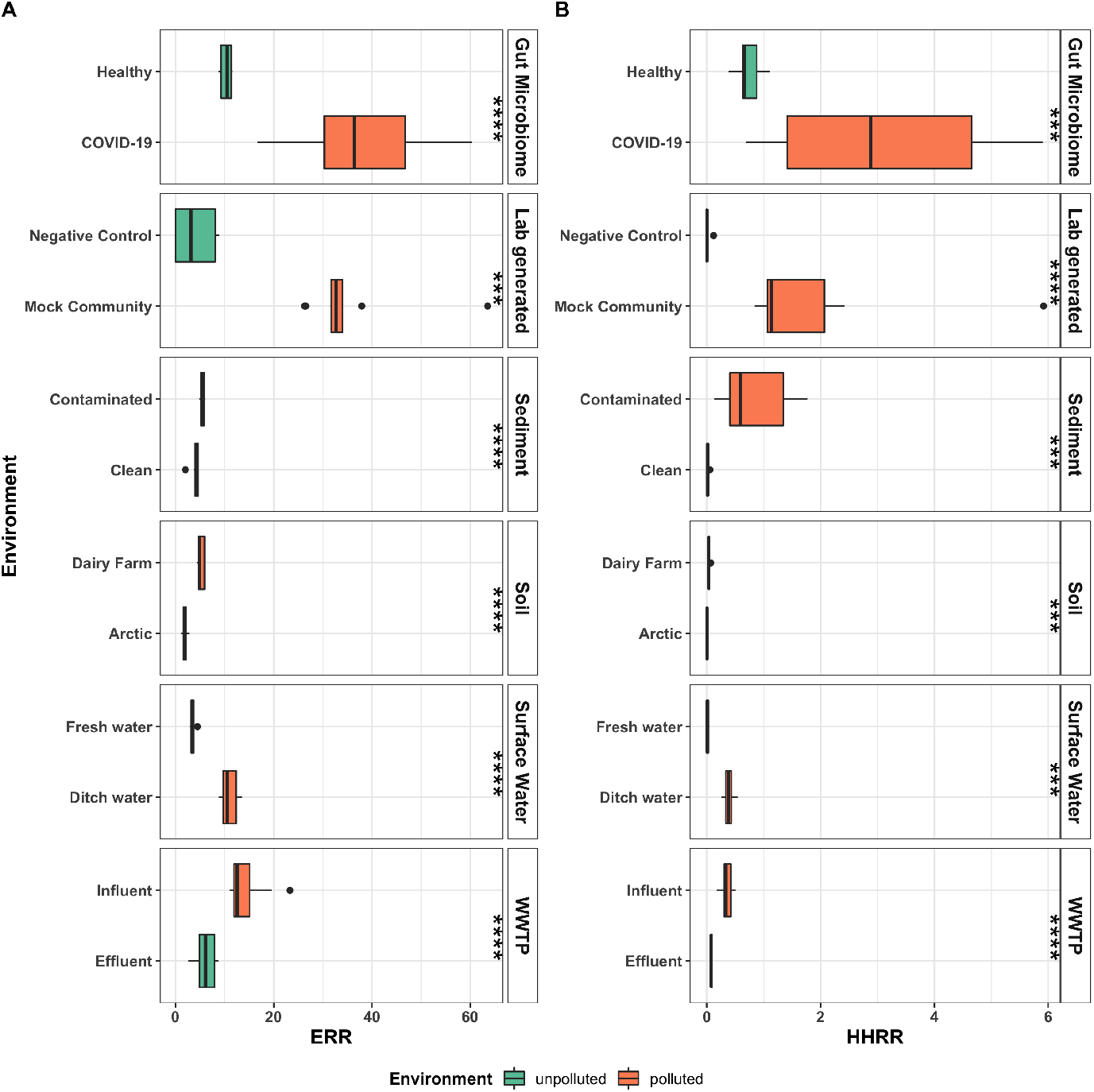
Range of ERR (A) and HHRR(B) for six different environments. Here, “polluted”/ “unpolluted” were generally defined to indicate higher/lower human contamination or other known impact and thus expected to be more enriched with ARGs, MGEs and pathogens. ^*^ p*<* 0.05; ^**^ p*<* 0.01, ^***^ p*<* 0.001, ^****^ p*<* 0.0001 according to Wilcoxon rank-sum test comparing scores between the paired categories.

**Fig. 4.**
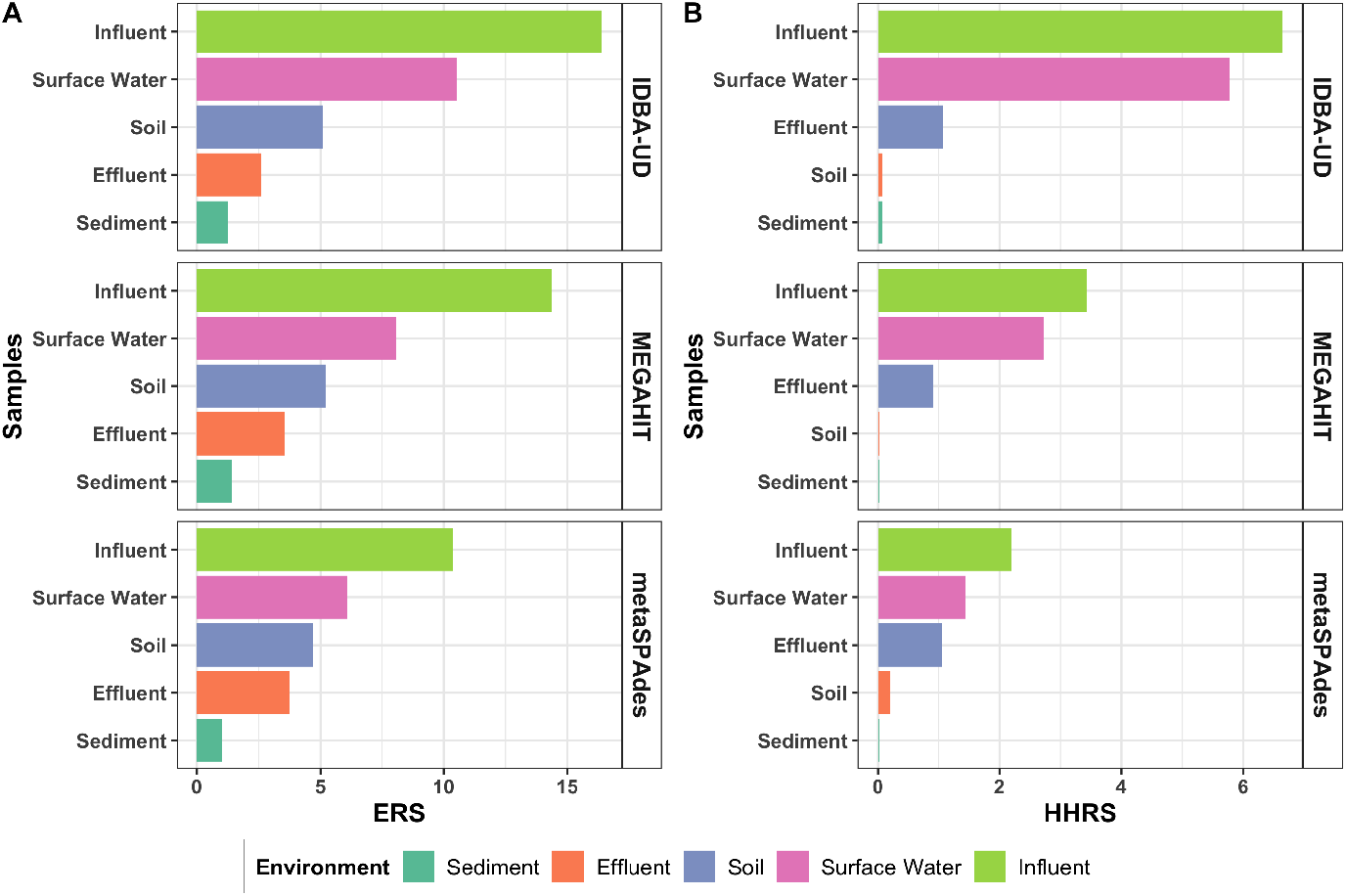
Effect of the assembler on the resistome risk score. Ranking of the resistome risk scores of the samples remain the same irrespective of the assembly method used.

### 4.2 Effect of Different Assembly Programs

The effect of assembly methodology on resistome risk score was tested by applying the MetaCompare 2.0 pipeline on contig sets obtained from the same samples using three different assemblers. This test was performed on influent wastewater, effluent wastewater, polluted soil, sediment, and surface water samples. Although the resistome risk scores differed for a given sample as a function of the assembler used to generate the contigs, the same general trend in the ranks of the resulting resistome risk scores was produced 4. This analysis indicates that overall trends in resistome risk scores are likely to hold true, even if different assemblers are applied. However, because the scores themselves differ, we recommend that users make resistome risk score comparisons among samples computed from contigs generated by the same assemblers.

### 4.3 Effect of Sequencing Depth and coverage

Given the complexity of environmental metagenomes, we further assessed the effect of sequencing depth and coverage on resistome risk scores and estimated the “saturation point” for converging scores. Simulated samples containing short reads at different depths of 1M, 10M, 50M, 100M, 125M, and 250M reads were assembled and then run through the pipeline. Figure 5(A and B) demonstrates that, although the ERR and HHRR scores vary with sequencing depth for the same environment, MetaCompare 2.0 was able to distinguish and produce the correct ranking of resistome risks among influent, effluent, and activated sludge environments over the broad range of sequencing depths. As expected, risk scores tend to vary more with low sequencing depth, e.g., 1M, 10M with coverage *≥* 50%, but become stable at 50M depth and *≤*60% coverage. This suggests that samples of similar sequencing depths are still comparable via MetaCompare 2.0, even with relatively shallow sequencing of 10M (Figure 5C). However, to be able to compare metagenomes of different depths, they should ideally be sequenced at *>* 60% coverage.

**Fig. 5.**
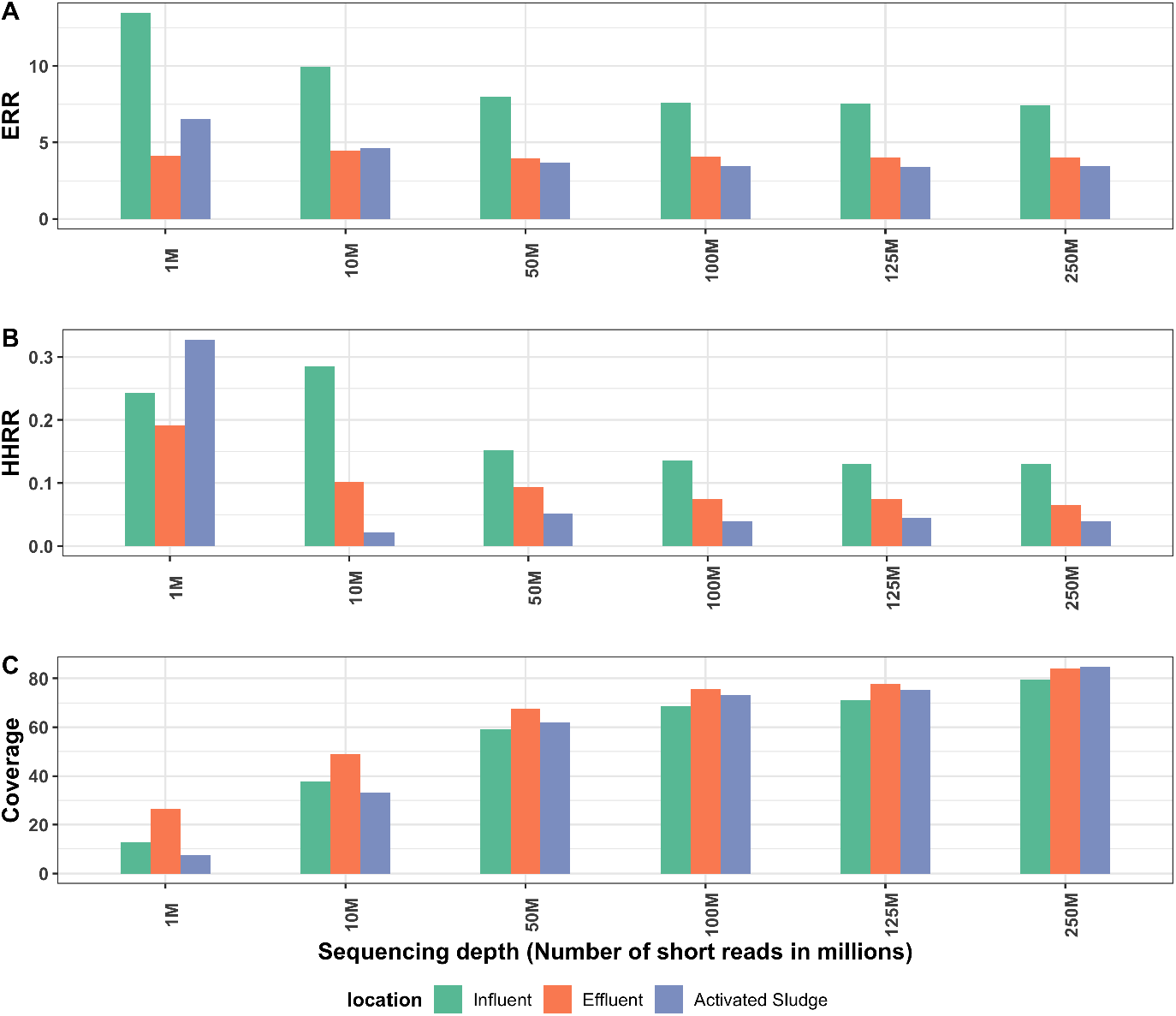
Effect of sequencing depth and library coverage on resistome risk scores of a subset of activated sludge samples. A) ERR for samples with different numbers of short reads. B) HHRR for samples with different numbers of short reads. C) Coverage of all samples in panel A/B. Risk scores reach a saturation level at 50M depth and *≥* 60%coverage.

Noticing high variability of scores generated using MetaCompare 1.0 among air samples reported in a recent study [31], we ran another experiment where multiple samples containing different numbers of contigs were generated from the largest activated sludge sample assembled in the previous experiment. We generated 50 sub-samples for each n, where n is the number of contigs. Figure 6(A and B)shows that the range of scores can trend higher for samples generating fewer contings (*≥* 10*k*) and the scores converge for samples with a higher number of contigs (*≤* 50*k*).

**Fig. 6.**
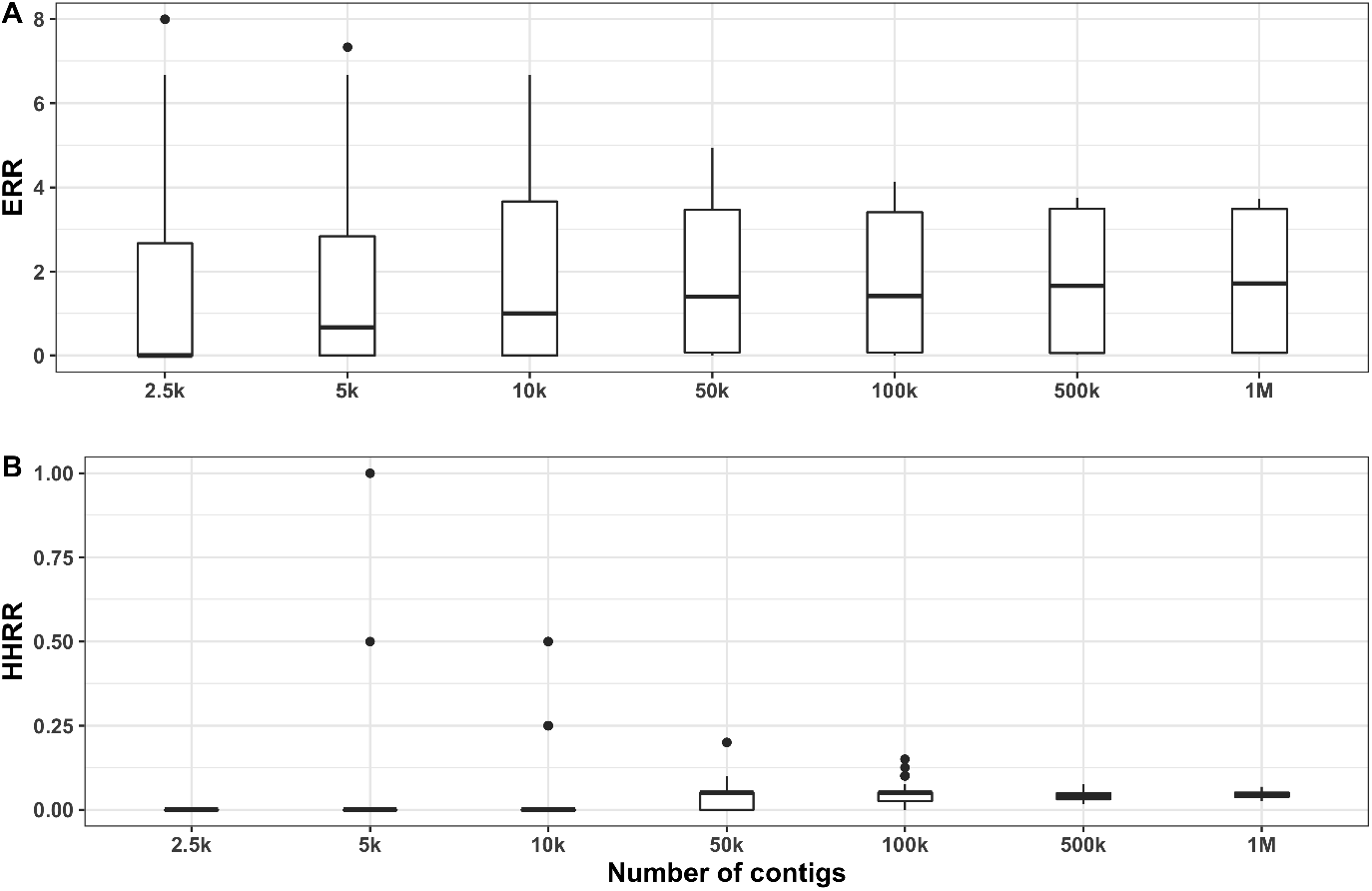
Range of risk scores for samples having different sizes of contig sets, A) ERR, B)HHRR. A higher variation of risk scores is observed for smaller (*<* 50*k*) contig sets.

## 5 Discussion

MetaCompare 2.0 provides several improvements over MetaCompare 1.0, including a revamped algorithm, faster annotation tools, updated databases, and a publicly-available and user-friendly interface. The determination of two distinct risk scores, ERR and HHRR, provides greater resolution in comparing the relative resistome risks across various environments of interest. The HHRR score is more closely aligned with conventional microbial risk assessment methodologies in a way that it focuses more on specific pathogens of concern (i.e., ESKAPE pathogens) and the acute human health hazards that they represent. The ERR, on the other hand, can be applied in scenarios where the concern is much broader in terms of assessing the potential for antibiotic resistance to evolve and spread, e.g., during microbial treatment of industrial wastewater or application of biosolids to soil. In sum, both the ERR and HRR can provide a comparative metric across a system, environment, or environments of interest to identify potential “hot-spots” worthy of additional attention from a mitigation standpoint. MetaCompare can also be used to assess trends with time, e.g., in response to a targeted intervention.

We tested MetaCompare 2.0 over a wide range of sample types in order to characterize the distribution of risk scores encountered. Consistent with expectation, sample sets with greater anthropogenic contamination/impact (i.e., “pollution”) yielded correspondingly higher resistome risk scores. We noticed higher risk scores in aquatic environments (e.g., wastewater, surface water) compared to terrestrial environments (e.g., soil, sediment). Interestingly, the gut microbiome samples exhibited the highest risk scores. Even the resistome risk scores for healthy humans produced higher scores than raw Influent wastewater (i.e., sewage) samples. This is consistent with the well-established understanding that human feces is enriched in pathogens, while here we see that these pathogen markers essentially become diluted with other microbes in the sewage collection network [53]. According to Zhang Z. et al. [17], human-associated environments (e.g., skin, gut) tend to have higher ARG abundances compared to other environments. A cross-environmental study can provide more insights into the composition of ARGs, MGEs, pathogens and their association with risk scores in such environments.

While we established a theoretical range of resistome risk scores between 0-100, empirically the general range of ERR scores was observed to be *<* 40 for “polluted” samples, and *<* 10 for “unpolluted” samples. The corresponding range of HHRR scores would be *<* 10 and ≈ 0, respectively. Such thresholds may prove helpful in identifying potential hotspots of concern for the dissemination of antibiotic resistance that warrant further attention in terms of further assessment or mitigation efforts.

In recent work, there have been multiple attempts in ranking risks associated with environmental aspects antibiotic resistance. Even though similar aspects have been considered in corresponding risk score computation, such as the mobility and pathogenicity of ARGs, there remains an important distinction between the MetaCompare framework and that proposed by others, that is, MetaCompare computes a “resistome risk” score for the full range of ARGs identified across a metagenome, whereas other approaches [16, 17] assign a risk score or rank for individual ARGs. Both frameworks have pros and cons, depending on the aim of the application. If one wants to compare samples derived from different environments in terms of aggregate risk of antibiotic resistance evolution and spread, having a single metric that captures the full range of ARGs is more relevant than focusing in on individual ARGs. However, if one is interested in zooming in on a specific subset of ARGs of concern, then ranking and scoring individual ARGs may be more useful. Arguably, ARGs with high “risks” in a sample should be more of the focal point for evaluating the risk score of the sample than those with low “risks”. Towards incorporating this aspect into the MetaCompare framework, MetaCompare 2.0 introduced the HHRS calculation, focusing specifically on ARGs and pathogens of most acute human health concern in the context of antibiotic resistance.

We further evaluated the effect of assemblers on the risk score calculation. For that purpose, we employed three assemblers, namely IDBA-UD [49], MEGAHIT [43], and MetaSPAdes [50], all of which are widely used in publicly-available pipelines. We observed in our experiment that the final scores were different for the contig sets generated by different assemblers, but the ranking of risk scores of the samples remained the same (Figure 4). Considering the required computing resources, speed and the tendency to produce more accurate assemblies [41, 54], we used MEGAHIT as the default assembler for all other experiments in this project. Though users are free to choose any assembly pipeline based on their requirements and preferences, we recommend use of a single assembler when comparing and ranking risk scores of samples.

We assessed how differences in sequencing depth might affect the scores. Theoretically, MetaCompare 2.0 can be applied on any sample irrespective of coverage and sequencing depth. However, it is noted that the risk scores for samples with either lower coverage or sequencing depth tended to have higher variance (Figure 5). This result is not surprising due to the highly stochastic nature of the set of genes or sequences that happen to be generated in the sequencing experiment. In fact, low coverage or sequencing depth can adversely affect contig assembly and thus downstream analyses resulting from the assembly [55]. Based on our experiments, we suggest that applying our pipeline on samples with *≥* 60% coverage would be the best practice.

Finally, we note that our evaluation of MetaCompare 2.0 was limited to samples sequenced on Illumina platform. Although long read data such as nanopore sequences were not evaluated in this study, theoretically MetaCompare 2.0 can also be applied to such data. One point of caution may be higher sequencing error rates associated with long-read technologies [56]. In such cases, the downstream pipeline could be adversely affected in terms of accuracy of annotation and may generate false negatives, although the trends would be expected to still be consistent. Still, the lower coverage that is typical when long-read sequening is applied for shotgun metagenomics could also be an issue, as it would not likely be feasible to achieve *≥* 60% coverage recommended here. Further validation of MetaCompare 2.0 for long-read metagenomics and benchmarking to Illumina sequencing would be recommended. At present, we advise that comparison of risk scores should be conducted amongst samples generated by the same sequencing platform rather than cross platforms.

**Supplementary information. Table S1**: List of pathogens included used in ecological resistome risk calculation. **Table S2**: Rank 1 ARGs and their homologs used in human health resistome risk calculation. **Table S3**: Samples used to evaluate range of risk scores in different environments. **Table S4**: Samples used to evaluate the effect of Assembly methods.

## Acknowledgments

We thank James Stoll for helping with the webserver setup.

## Declarations

### Code availability

The web service for MetaCompare 2.0 is available at http://metacompare.cs.vt.edu/. The source code for the entire pipeline can be found at https://github.com/mrumi/MetaCompare2.0. Users are required to create an account before uploading sequences to the platform.

## Author’s Contributions

M.A. Rumi performed all the computations, carried out experiments, and wrote the manuscript with support from B.C. Davis, P.J. Vikesland, A. Pruden, and L. Zhang. M. Oh developed the theory and designed the website. A. Juvekar contributed to the design of the visualization module of the website. B.C. Davis and C.L. Brown helped develop the theory and experimental design. A. Pruden and P.J. Vikesland helped supervise the project. L. Zhang supervised the project.

## Ethics declarations

### Ethics approval and consent to participate

Not applicable

### Consent for publication

This document has been reviewed in accordance with U.S. Environmental Protection Agency policy and approved for publication. The views expressed in this article are those of the authors and do not necessarily represent the views or policies of the U.S. Environmental Protection Agency. Mention of trade names, products, or services does not convey, and should not be interpreted as conveying, official EPA approval, endorsement or recommendation.

